# A distinct signalling circuit upregulates cytotoxic ROS production in a cellular model of Parkinson’s disease: key roles for TRPM2, Zn^2+^ and complex III

**DOI:** 10.1101/2024.01.08.574664

**Authors:** Maali AlAhmad, Hala Isbea, Esra Shitaw, Fangfang Li, Asipu Sivaprasadarao

**Affiliations:** School of Biomedical Sciences, Faculty of Biological Sciences, University of Leeds, Leeds, LS29JT, UK; Department of Biological Sciences, College of Science, Kuwait University, Alshadadiya, PO Box 5969, Safat, 130602, Kuwait; Center for Rehabilitation Medicine, Rehabilitation & Sports Medicine Research Institute of Zhejiang Province, Department of Rehabilitation Medicine, Zhejiang Provincial People’s Hospital, Affiliated People’s Hospital, Hangzhou Medical College, Hangzhou,China

**Author notes:** Equal contribution (Maali AlAhmad, Hala Isbea). Corresponding author Professor Asipu Sivaprasadarao School of Biomedical Sciences G6.44d, Garstang Building University of Leeds, Leeds, LS2 9JT UK.

## Abstract

Reactive oxygen species (ROS) serve vital physiological functions, but aberrant ROS production contributes to numerous diseases. Unfortunately, therapeutic progress targeting pathogenic ROS has been hindered by limited understanding of whether the mechanisms driving pathogenic ROS differ from those governing physiological ROS generation. To address this knowledge gap, we utilised a cellular model of Parkinson’s disease (PD), as an exemplar of ROS-associated diseases. We exposed SH-SY5Y neuroblastoma cells to the PD-toxin, MPP^+^ (1-methyl-4-phenylpyridinium) and studied ROS upregulation leading to cell death, the primary cause of PD. We demonstrate: (i) MPP^+^ stimulates ROS production by raising cytoplasmic Ca^2+^ levels, rather than acting directly on mitochondria. (ii) To raise the Ca^2+^, MPP^+^ co-stimulates NADPH oxidase-2 (NOX2) and the TRPM2 (Transient Receptor Potential Melastatin2) channel that form a positive feedback loop to support each other’s function. (iii) Ca^2+^ exacerbates mitochondrial ROS (mtROS) production not directly, but via Zn^2+^. (iv) Zn^2+^ promotes electron escape from respiratory complexes, predominantly from complex III, to generate mtROS. These conclusions are drawn from data, wherein inhibition of TRPM2 and NOX2, chelation of Ca^2+^ and Zn^2+^, and prevention of electron escape from complexes -all abolished the ability of MPP^+^ to induce mtROS production and the associated cell death. Furthermore, calcium ionophore mimicked the effects of MPP^+^, while Zn^2+^ ionophore replicated the effects of both MPP^+^ and Ca^2+^. Thus, we unveil a previously unrecognized signalling circuit involving NOX2, TRPM2, Ca ^2+^, Zn^2+^, and complex III that drives cytotoxic ROS production. This circuit lies dormant in healthy cells but is triggered by pathogenic insults and could therefore represent a safe therapeutic target for PD and other ROS-linked diseases.

## INTRODUCTION

Reactive oxygen species (ROS) play vital roles in both physiological and pathological processes. In healthy eukaryotes, ROS generation is a continuous process, and cellular antioxidant defence mechanisms promptly neutralize any excess ROS ^1, 2, 3^. ROS, such as hydrogen peroxide (H_2_O_2_) and superoxide (O_2_^.-^), initiate reversible modifications of specific cysteine residues in kinases and phosphatases, thereby regulating various physiological processes including immunity, cell proliferation, development, and cognition ^3, 4^. However, in pathological conditions, the production of H_2_O_2_ and O_2_^.-^ increases, leading to their conversion into highly reactive hydroxyl radicals (^.^OH). These radicals cause irreversible and nonspecific modifications of proteins, lipids, and nucleic acids ^1, 2, 3^, resulting in pathogenic signalling that contributes to accelerated aging and diverse pathologies (>30), ranging from diabetes to cardiovascular diseases to neurodegenerative disorders such as Alzheimer’s and Parkinson’s diseases ^5, 6, 7^.

However, antioxidant supplements have yielded disappointing outcome in clinical trials ^2, 6, 8^. This lack of success could be attributed to the inability of antioxidants to discriminate pathogenic ROS from physiological ROS ^6, 7, 8, 9^. Furthermore, scavenging of pathogenic ^.^OH is thought to be impractical ^8^. Thus, to develop safe and effective redox medicines, it is imperative to gain a better understanding of how the shift from physiological to pathological ROS status occurs.

However, ROS biology is far too complex, involving multiple signals and mechanisms. One key signal is Ca^2+^ ^10, 11^. Indeed, there is a direct correlation between the cytosolic Ca^2+^ rise and the amount of ROS produced ^10, 11^. In healthy cells, physiological stimuli induce a modest rise in cytosolic Ca^2+^ to stimulate ROS production at levels appropriate for redox signalling. Pathogenic insults, however, rise the Ca^2+^ to supraphysiological concentrations to bolster ROS production to toxic levels ^10, 11^. ROS are produced from multiple sources, including mitochondrial complexes and dehydrogenases, and extra-mitochondrial enzymes, such as NADPH oxidases (NOXs) ^1, 2, 6, 8, 10, 12^. Furthermore, ROS produced at one site can impact ROS generation at a different site- a phenomenon known as ROS-induced ROS production ^12, 13^, which adds an additional layer of complexity to ROS homeostasis.

Regulation of Ca^2+^ homeostasis is equally complex, involving various ion channels, exchangers and transporters that transport Ca^2+^ between the Ca^2+^ rich compartments and cytoplasm ^10, 11^. The picture gets even more complex by the existence of a Ca^2+^-ROS feedback cycle, where ROS can trigger cytosolic Ca^2+^ rise and Ca^2+^, in turn, can stimulate ROS production^10, 11, 12^. The precise mechanisms underlying the Ca^2+^-ROS cycle are not fully understood but appear to involve an intricate interplay between ROS-sensitive calcium channels and Ca^2+^-dependent ROS-generation from NADPH oxidases (NOXs) and mitochondria ^10^.

Some Ca^2+^ channels are ROS sensitive and therefore can contribute to Ca^2+^-ROS interplay ^10^. Among them, TRPM2 calcium channels are important. These channels are co-activated by ROS-generated ADPR and Ca^2+^ ^14^ ^15^. Importantly, activation of these channels has been linked to numerous diseases where ROS are upregulated ^16, 17^. Therefore, we hypothesise that TRPM2 channels play a central role in Ca^2+^-ROS interplay.

To test this hypothesis, we utilized a well-established cellular model of Parkinson’s disease (PD) ^18^ as an exemplar of ROS-linked diseases. We exposed SH-SY5Y neuroblastoma cells to the PD-causing neurotoxin MPP^+^ (1-methyl-4-phenylpyridinium) to upregulate Ca^2+^ ^19^ and investigated how the rise in Ca^2+^ augments ROS production leading to cell death. Our data reveal a distinct signalling circuit-comprising NOX2, the TRPM2 channel, Ca^2+^, Zn^2+^, and respiratory complexes I and III, predominantly the latter- that links Ca^2+^ to pathogenic levels of ROS production.

## RESULTS

### Amplification of mitochondrial ROS is essential for the neurotoxic effect of MPP^+^

We first investigated the relative contributions of mitochondrial and extra-mitochondrial sources to MPP^+^-induced cell death. For this, we exposed SH-SY5Y cells to 1 mM MPP^+^ for 24 hours and assessed ROS production and cell viability (for optimisation of conditions, see Supplemental Figure 1). We used dihydroethidium (DHE) to detect total ROS from all cellular sources, including mitochondria, and MitoSOX™ Red to specifically detect mtROS. MPP ^+^ treatment significantly increased the fluorescence of both the reporters. The total ROS signal was abolished by the general ROS quenchers, N-acetylcysteine (NAC) and TEMPO (Figure 1 A-B), confirming that the increase in fluorescence is due to ROS. Interestingly, Mito-TEMPO, a ROS quencher specific for mtROS, was able to abolish the total ROS signal almost as effectively as the general ROS quenchers (Figure 1 A-C). Furthermore, Mito-TEMPO was able to prevent MPP^+^-induced cell death (Figure 1 D-E). These results are seemingly in agreement with the numerous reports that MPP^+^ acts directly on mitochondria to produce cytotoxic ROS ^20^.

**Figure 1:**
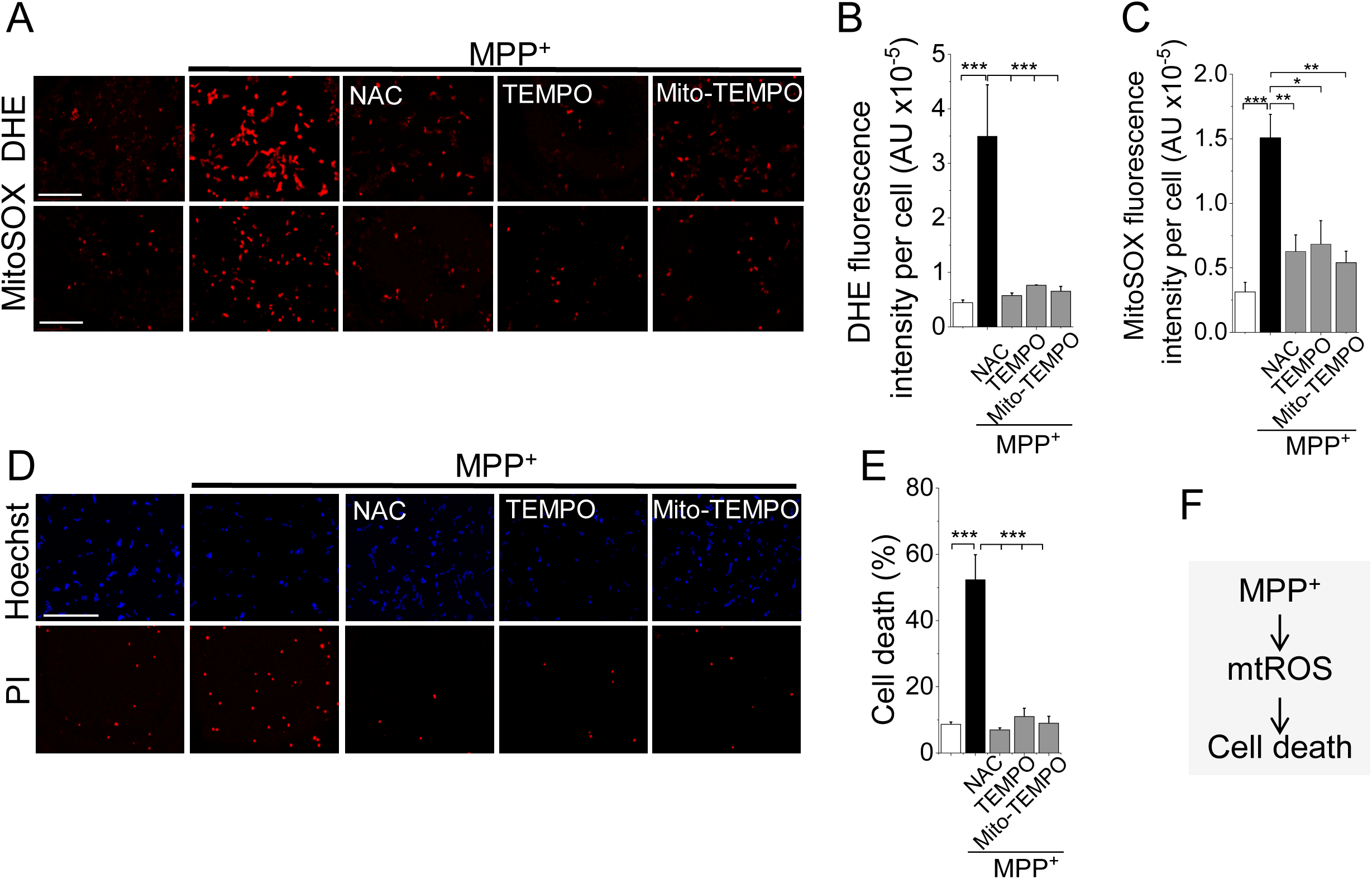
MPP^+^ exacerbates mitochondrial ROS production to cause SH-SY5Y cell death. **(A-C)** MPP^+^ induces ROS production. **(A)** Fluorescent images of SH-SY5Y cells stained for total (DHE) and mitochondrial (MitoSOX) ROS, recorded after 24-hour treatment with medium alone or medium containing 1 mM MPP^+^ ± the indicated antioxidants (5 mM NAC, 10 µM TEMPO or 10 µM Mito-TEMPO); scale bar =75 µm. **(B-C)** Mean ± SEM of fluorescence intensity corresponding to (A). **(D-E)** MPP^+^ induces cell death. **(D)** Fluorescent images of cells treated as in (A), but stained for cell death (Hoechst and PI). **(E)** Mean ± SEM of % PI positive dead cells corresponding to (D); scale bar= 125 µm. Mean data in B, C and E are from three independent experiments (n =3); * *p* ˂ 0.05, ** *p* ˂ 0.01; *** *p* ˂ 0.001 from one-way Anova, post-hoc Tukey test. (**F**) Schematic summary of findings.

### MPP^+^ upregulates mitochondrial ROS production *indirectly* by generating Ca^2+^ and Zn^2+^ signals

It is widely believed that MPP^+^ acts directly on respiratory complex I to generate mtROS^20^ ^21^. However, MPP^+^ has also been shown to elevate cytoplasmic Ca^2+^ ^19^. Since Ca^2+^ is a known stimulant of mtROS^10, 11^, we asked whether MPP^+^-induced mtROS production could, in part, be Ca^2+^ mediated. As expected, MPP^+^ treatment caused an increase in intracellular Ca^2+^ levels (Figure 2A-B). Rather unexpectedly, chelation of Ca^2+^ with BAPTA completely abolished the ability of MPP^+^ to stimulate mtROS production (Figure 2C-E). These data suggested that the MPP^+^-induced mtROS production is almost entirely mediated by Ca^2+^, and/or other metal ions that BAPTA is capable of binding. We considered Zn^2+^ as the other metal ion because its affinity for BAPTA is greater than that for Ca^2+^, and Zn^2+^ is a known stimulant mtROS production ^22^. Thus, we tested the effect of the Zn^2+^ chelator, TPEN (N,N,N,’,N’-tetrakis (2-pyridylmethyl) ethylenediamine), using a low concentration to chelate Zn^2+^, while excluding Ca^2+^ ^23^. Remarkably, TPEN was as effective as BAPTA in suppressing MPP^+^-induced ROS production (Figure 2C-E) and the consequent cell death (Figure 2F-G). The ability of the two chelators to completely abolish MPP^+^-induced ROS production argues against the numerous reports that MPP^+^ acts directly and solely on mitochondria ^24, 25^. Instead, our results suggest that Ca^2+^ depends on Zn^2+^ signals to upregulate ROS to cytotoxic levels.

**Figure 2:**
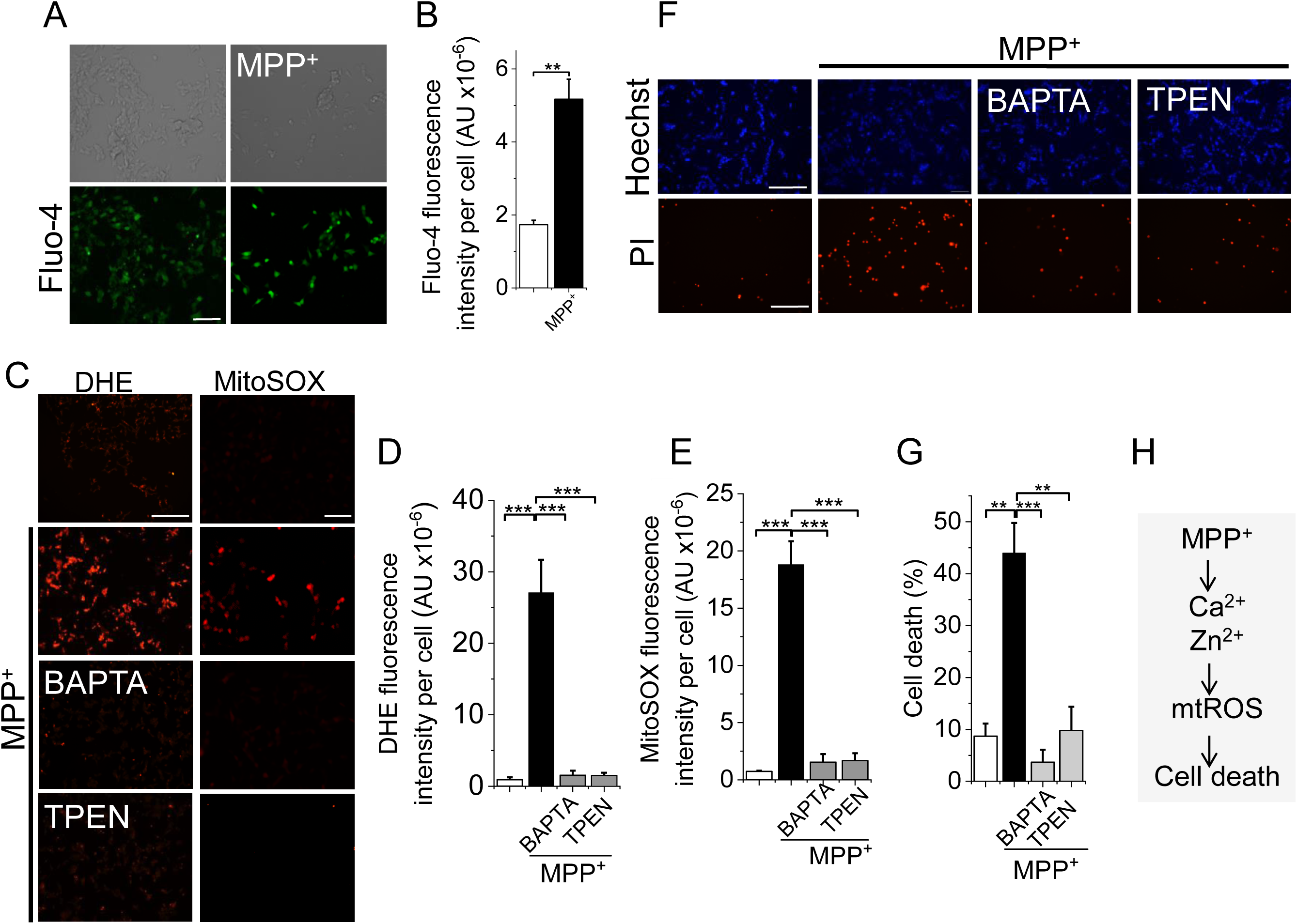
MPP^+^ upregulates intracellular Ca^2+^ and Zn^2+^ but chelation of Zn^2+^ alone is sufficient to prevent mtROS overproduction and cell death. (**A-B**) MPP^+^ causes a rise in intracellular Ca^2+^. (**A**) Fluorescent images of SH-SY5Y cells stained for Ca^2+^ with Fluo-4 imaged after 24 hour treatment with medium alone or medium containing 1 mM MPP^+^; scale bar = 75 µm. (**B**) Mean ± SEM of fluorescence intensity. (**C-E**) Chelation of Zn^2+^ is as effective as Ca^2+^ chelation in attenuating ROS production. (**C**) Fluorescent images of cells stained for total (DHE) and mitochondrial (MitoSOX) ROS, imaged after 24 hour treatment with medium alone or medium containing 1 mM MPP^+^ ± BAPTA-AM (5 µM) or TPEN (0.5 µM); scale bar = 75 µm. (**D-E**) Mean ± SEM of fluorescence intensity. (**F-G**) Chelation of Zn^2+^ is as effective as Ca^2+^ chelation in preventing MPP^+^-induced cell death. (**F**) Fluorescent images of cells treated as in (C) but stained for cell death (Hoechst and PI); scale bar = 125 µm. (**G**) Mean ± SEM of % PI positive dead cells. Mean data in B, D, E and G are from three independent experiments (n =3); ** *p* ˂ 0.01; *** *p* ˂ 0.001 from One-way Anova, post-hoc Tukey test. (**H**) Schematic summary of findings.

### TRPM2-mediated Ca^2+^ rise upregulates mitochondrial ROS to cytotoxic levels

We next set out to investigate the mechanism by which MPP^+^ raises cytoplasmic Ca^2+^ levels. A previous study has shown that MPP^+^ can stimulate Ca^2+^ entry via TRPM2 channels ^19^. However, neuronal cells have multiple mechanisms to regulate Ca ^2+^ ^26^. Thus, it was not clear whether TRPM2-mediated Ca^2+^ entry alone can account for MPP^+^ induced ROS production. Accordingly, we tested the effect of inhibiting TRPM2-mediated Ca^2+^ influx on ROS production. Inhibition of TRPM2 channels with three pharmacological inhibitors (PJ34, 2-APB, and ACA) with different modes of action ^27^ abolished MPP^+^-induced mtROS production (Figure 3A-C) and cell death (Figure 3D-E). Likewise, silencing RNA targeted to the TRPM2 channel, but not the control scrambled siRNA, abrogated ROS production (Figure 3F-H) and cell death (Figure 3I-J). Importantly, there was no significant difference between control cells and cells exposed to MPP^+^ but had been pre-treated with TRPM2 inhibitors/siRNA (Figure 3B-C, G-H). These results imply that TRPM2 mediated Ca^2+^ entry fully accounts for the MPP^+^-induced ROS production and has little or no effect on basal ROS production.

**Figure 3:**
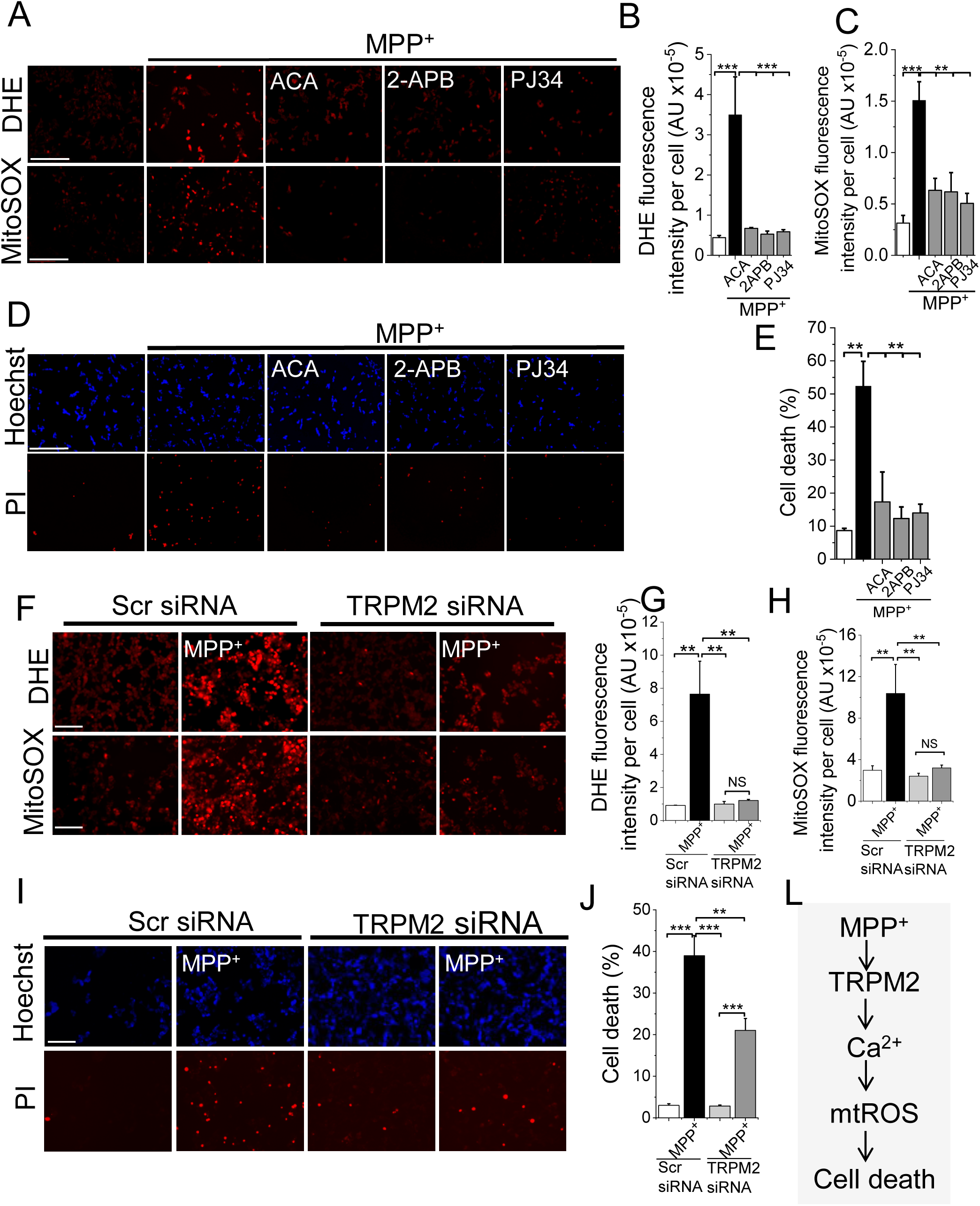
Inhibition of TRPM2 calcium channels prevents MPP^+^ induced ROS production and SH-SY5Y cell death. (**A-E**) Pharmacological inhibition of TRPM2 channels prevents MPP^+^-induced ROS generation and SH-SY5Y cell death. Cells were treated with medium alone or medium containing 1 mM MPP^+^ ± TRPM2 inhibitors: ACA (10 µM), 2-APB (50 µM) and PJ34 (10 µM). (**A**) Fluorescent images of cells stained for total (DHE) and mtROS (Mito-SOX); scale bar = 125 µm. (**B-C**) Mean ± SEM of fluorescence intensity. (**D**) Fluorescent images of cells treated stained for cell death (Hoechst and PI); scale bar = 125 µm. (**E**) Mean ± SEM of % PI positive dead cells. (**F-J**) TRPM2-targeted siRNA attenuates MPP^+^-induced ROS generation and SH-SY5Y cell death. Cells transfected with TRPM2 siRNA or scrambled (Scr) siRNA were treated with medium or medium containing MPP^+^. (**F**) Fluorescent images of cells stained for total (DHE) and mtROS (Mito-SOX); scale bar = 125 µm. (**G-H**) Mean ± SEM of fluorescence intensity. (**I**) Fluorescent images of cells stained for cell death (Hoechst and PI); scale bar = 125 µm. (**J**) Mean ± SEM of % PI positive dead cells. Mean data in B, C, E, G, H and J are from three independent experiments (n =3); ** *p* ˂ 0.01; *** *p* ˂ 0.001 from One-way Anova, post-hoc Tukey test. (**K**) Schematic summary of the findings.

To seek independent evidence for the exclusive role of TRPM2 in mtROS generation, we used HEK-293 cells stably transformed with the FLAG-tagged human TRPM2 cDNA placed under the control of a tetracycline-regulated promoter (HEK-TRPM2^tet^) ^28, 29^. Western blotting confirmed the expression of TRPM2 channels in tetracycline induced, but not in non-induced, cells (Figure 4A). Functional analysis showed that H_2_O_2_ was able to induce robust Ca^2+^ influx in the induced, but not non-induced, cells (Figure 4B). Using these validated cells, we demonstrated that H_2_O_2_ had virtually no effect on ROS production in uninduced cells, but triggered robust mtROS production in induced cells that was sensitive to 2-APB inhibition (Figure 4C-E).

**Figure 4:**
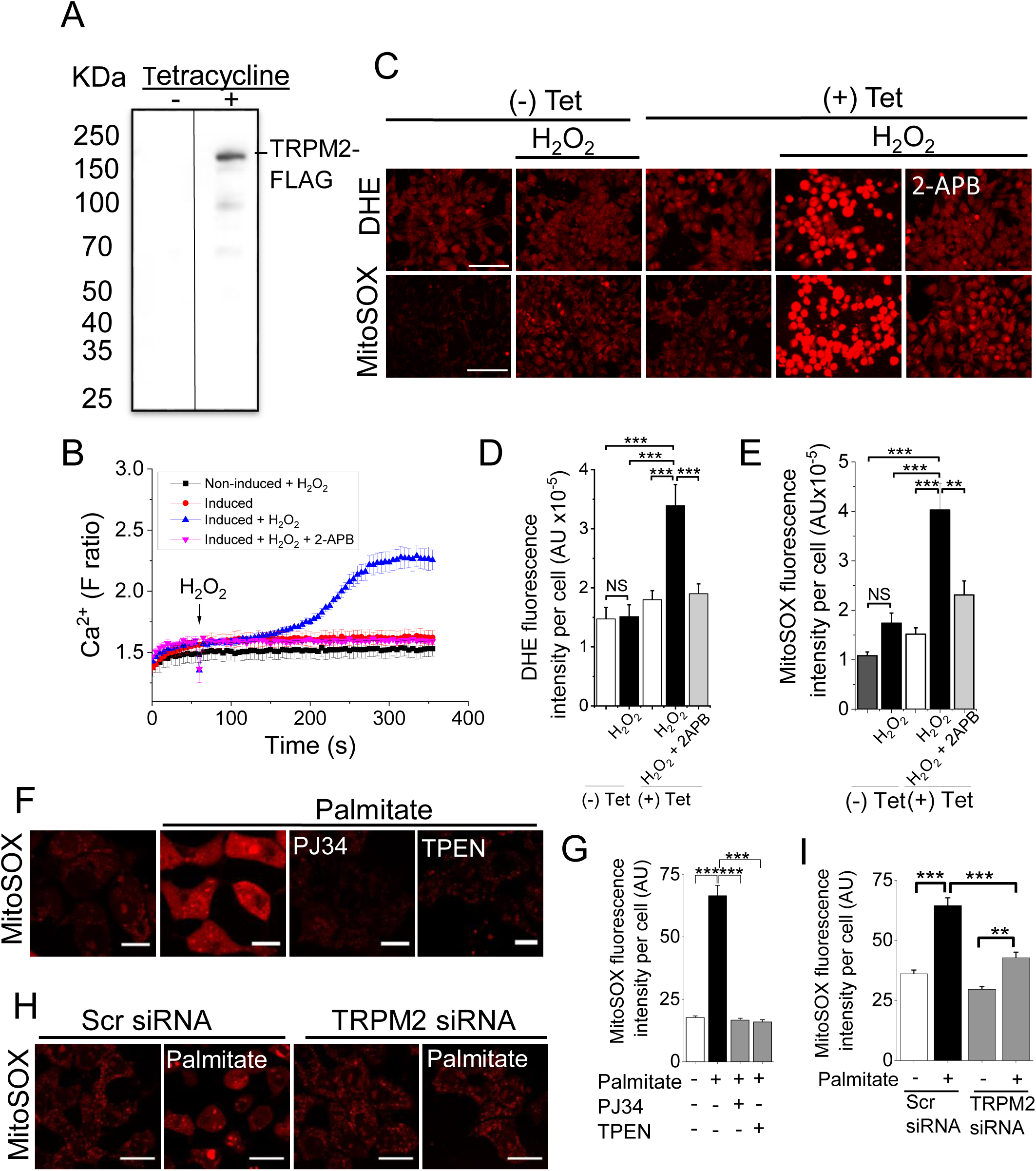
Activation of TRPM2 channels is essential for stress-induced ROS production in other cell lines. **(A-B)** Tetracycline induction of TRPM2-FLAG expression in HEK-TRPM2^tet^ cells. **(A)** Western blot shows a band corresponding to the size of TRPM2-FLAG in lysates of induced, but not uninduced HEK-TRPM2^tet^ cells. **(B)** Tetracycline-induced HEK-TRPM2^tet^ cells, but not uninduced or 2-APB-treated cells, show H_2_O_2_-induced Ca^2+^ rise (blue trace), monitored with Fura-2. **(C-E)** TRPM2 expression is essential for mtROS overproduction. Tetracycline induced and uninduced HEK-TRPM2^tet^ cells were treated with medium alone or medium containing H_2_O_2_ (50 µM) ± 2-APB (50 µM) for 2 hours before staining for ROS. (**C**) Fluorescent images of cells stained for total (DHE) and mitochondrial (Mito-SOX) ROS; scale bar = 75 µm. (**D-E**) Mean ± SEM of fluorescence intensity. (**F-I**) Palmitate-induced mtROS production in INS1-832/13 pancreatic β-cell line depends on TRPM2 function and Zn^2+^. (**F-G**) TRPM2 inhibition and Zn^2+^ chelation attenuates palmitate-induced mtROS production. INS1-832/13 cells were treated with medium containing the vehicle (human serum albumin, HSA) or 500 µM palmitate - HSA complex in the absence or presence of PJ34 (10 µM) or TPEN (1 µM) for 4 hours. (**F**) Confocal images of cells stained with MitoSOX; scale bar = 5 µm (**G**) Mean ± SEM of fluorescence intensity. (**H-I**) TRPM2-targeted siRNA attenuates palmitate-induced mtROS production. Cells were transfected with scrambled siRNA (Scr siRNA) or TRPM2 siRNA before exposing to palmitate. (**H**) Confocal images of cells stained with MitoSOX; scale bar =10 µm (**I**) Mean ± SEM of fluorescence intensity. Mean data in D, E, G and I are from three independent experiments (n =3); ** *p* ˂ 0.01; *** *p* ˂ 0.001 from One-way Anova, post-hoc Tukey test.

Finally, we used a non-neuronal cell line, the pancreatic β-cell line (INS-1 832/13) that natively expresses TRPM2 channels. These cells are sensitive to palmitate induced nutrient stress and have been used as a cellular model for obesity-induced type-2 diabetes^29^. The results show rescue of palmitate-induced mtROS production by both the pharmacological inhibitor PJ34 (Figure 4F-G) and TRPM2 siRNA (Figure 4H-I). In addition, TPEN attenuated palmitate-induced mtROS production, indicating that the role of Zn^2+^ is not limited to neuronal cells (Figure 4F-G).

Thus, using three different cellular models and three different stressors, we demonstrated that TRPM2-mediated Ca^2+^ entry upregulates mtROS production to cytotoxic levels, but has little or no impact on basal ROS production. Thus, TRPM2 dependent pathogenic ROS production appears to be not unique to neuronal cells but shared by other cell types. Co-incidentally, the absolute dependence of MPP^+^ on TRPM2 channels for mtROS overproduction further excludes the direct action of MPP^+^ on mitochondria.

### TRPM2-mediated Ca^2+^ influx requires simultaneous activation of NADPH oxidase-2

TRPM2 channels require ROS for activation. Given the above evidence that MPP^+^ cannot directly stimulate ROS from mitochondria (Figures 2-4), we asked whether MPP^+^ could target NOX2 ^30^ to generate the ROS required for TRPM2 activation. Consistent with this idea, inhibition of NOX2 with apocynin (a pan NOX inhibitor) and gp91 -ds-tat (a NOX-2-specific inhibitor) prevented MPP^+^-induced Ca^2+^ rise (Figure 5A-B), and the consequent ROS production (Figure 5C-E) and cell death (Figure 5F-G). These findings support the idea that NOX2 activation likely provides the ROS necessary for TRPM2 activation and the subsequent mtROS upregulation and cell death.

Taken together, these results suggest that simultaneous activation of NOX2 and TRPM2 channels is necessary for MPP^+^ to generate the Ca^2+^ signals required for mtROS production and cell death.

**Figure 5:**
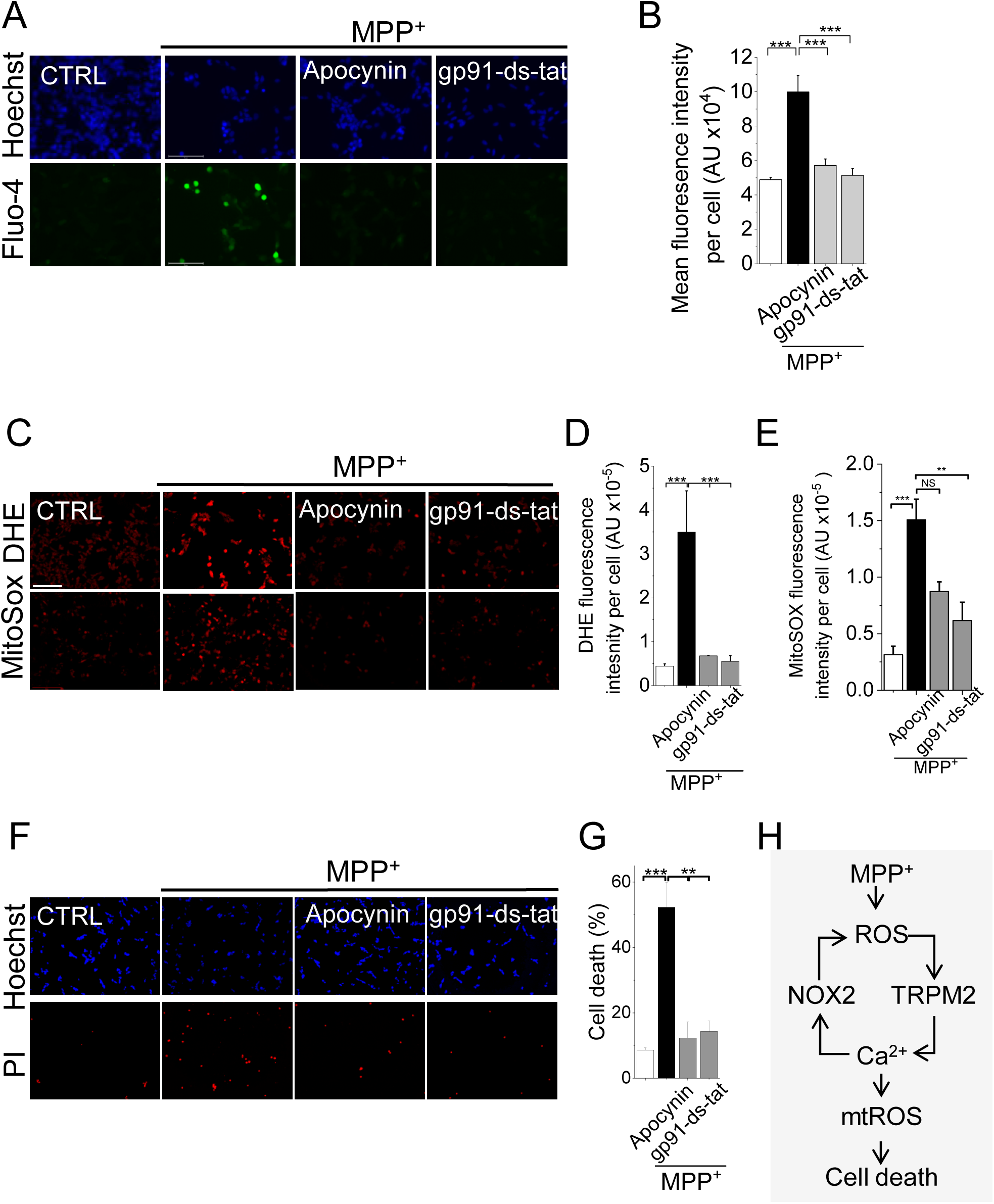
NOX2 inhibition prevents MPP^+^-induced intracellular Ca^2+^ rise, mitochondrial ROS production and cell death. (**A-B**) NOX2 inhibition prevents MPP^+^-induced Ca^2+^ rise. SH-SY5Y cells were treated with medium alone, or medium containing 1 mM MPP^+^ ± NOX inhibitors (10 µM apocynin and 5 µM Gp91ds-tat) for 24 hours. (**A**) Fluorescent images of cells stained for Ca^2+^ with Fluo-4; scale bar = 75 µm. (**B**) Mean ± SEM of fluorescence intensity. **(C-E)** NOX2 inhibition prevents MPP^+^-induced increase in total as well as mitochondrial ROS production. Cells were treated as in (A) and stained for ROS. (**C**) Fluorescent images of cells stained for total (DHE) and mtROS (Mito-SOX); scale bar = 125 µm. (**D-E**) Mean ± SEM of fluorescence intensity. **(F-G)** NOX2 inhibition abolishes MPP^+^-induced cell death. (**F**) Fluorescent images of cells treated as in (A) but stained for cell death (Hoechst and PI). **(G)** Mean ± SEM of % PI positive dead cells. Mean data in B, D, E, and G are from three independent experiments (n =3); ** *p* ˂ 0.01; *** *p* ˂ 0.001 from One-way Anova, post-hoc Tukey test. (**H**) Schematic summary of the findings.

### Zn^2+^ acts downstream of Ca^2+^ to augment mitochondrial ROS production to cytotoxic levels

While the above data (Figures 2-4) are entirely consistent with the established role that Ca^2+^ plays a role in mtROS production, they fail to explain why the Zn^2+^ chelator was as effective as the Ca^2+^ chelator in suppressing mtROS production (Figure 2). To address this, we postulated a hierarchical relationship between the two cations, with Zn^2+^ acting downstream of Ca^2+^.

To test the hypothesis, we raised the intracellular Ca^2+^ by treating the cells with the calcium ionophore A23187. We have included the membrane impermeable Zn^2+^ chelator DTPA (diethylenetriaminepentaacetic acid) during the treatment to exclude unwarranted Zn^2+^ entry. A23187 promoted cytosolic Ca^2+^ rise (Figure 6A-B), increasing the ROS production (Figure 6C-E) and cell death (Figure 6F-G). However, TPEN was able to mitigate all these effects as effectively as BAPTA (Figure 6C-G). These data support the hypothesis that Zn^2+^ acts downstream of Ca^2+^ to upregulate cytotoxic mtROS production.

**Figure 6:**
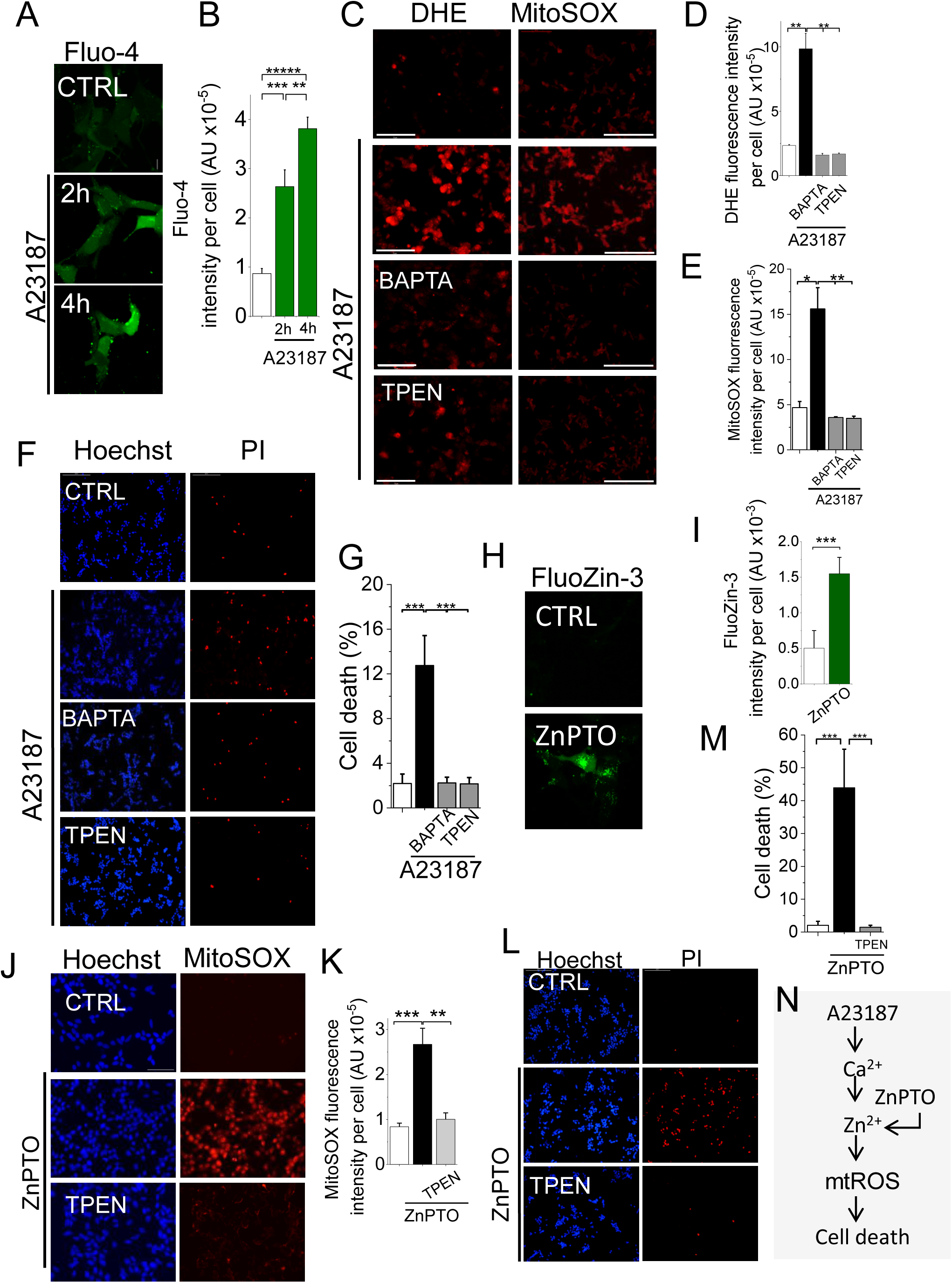
Zn^2+^ is essential for Ca^2+^-induced mtROS production. (**A-F**) A23187 (calcium ionophore)-mediated elevation of intracellular Ca^2+^ increases mtROS production and SH-SY5Y cell death, but Zn^2+^ chelation attenuates these effects. (**A-B**) A23187 mediated rise in intracellular Ca^2+^. Cells were treated with vehicle or A23187 (1 µM plus 1 mM DTPA) and stained with Fluo-4. (**A**) Representative confocal images of Fluo-4-stained cells; scale bar 14 µm. (**B**) Mean ± SEM of fluorescence intensity. (**C-G**) Chelation of both Ca^2+^ and Zn^2+^ prevented A23187-induced mtROS production and cell death. SH-SY5Y cells were treated with vehicle or A23187 (1 µM, plus 1 mM DTPA) with and without BAPTA -AM (5 µM) or TPEN (0.5 µM). After 4 hours incubation at 37°C, cells were stained for ROS and cell death. (**C**) Fluorescent images of cells stained for total (DHE) and mtROS (Mito-SOX); scale bar = 125 µm. (**D-E**) Mean ± SEM of fluorescence intensity. (**F**) Fluorescent images of cells stained for cell death (Hoechst and PI); scale bar =125 µm. (**G**) Mean ± SEM of % PI positive dead cells. (**H-K**) Elevation of intracellular Zn^2+^ with zinc ionophore (ZnPTO) induces mtROS production and cell death. SH-SY5Y cells were treated with vehicle (DMSO) or ZnPTO (2 µM, 2 hr, 37°C) and then stained for Zn^2+^ (FluoZin-3) or mtROS or cell death. (**H-I**) ZnPTO elevates intracellular Zn^2+^. (**H**) Representative confocal images of FluoZin-stained cells; scale bar = 14 µm. (**I**) Mean ± SEM data of fluorescence intensity. (**J-K**) ZnPTO induces mtROS production. (**J**) Fluorescent images of cells stained for mtROS (Mito-SOX); scale bar = 125 µm. (**K**) Mean ± SEM of fluorescence intensity. (**L-M**) ZnPTO induces cell death. (**L**) Fluorescent images of cells stained for cell death (Hoechst and PI); scale bar =125 µm. (**M**) Mean ± SEM of % PI positive dead cells. Mean data in B, D, E, G, I, K and M are from three independent experiments (n =3); * *p* ˂ 0.05**; *p* ˂ 0.01; *** *p* ˂ 0.001 from One-way Anova, post-hoc Tukey test. (**N**) Schematic summary of the findings.

To confirm that Zn^2+^ is the ultimate ROS-inducing signal, we elevated the intracellular Zn^2+^ levels with the zinc ionophore, zinc-pyrithione (ZnPTO). Ca^2+^ was excluded from the treatment medium to avoid any confounding effects from Ca^2+^ entry. ZnPTO treatment caused a rise in intracellular free Zn^2+^ (Figure 6H-I) and has increased mtROS production (Figure 6J-K) and cell death (Figure 6L-M), the effects being rescued by TPEN (Figure 6J-M). Thus, raising the intracellular Zn^2+^ was able to faithfully phenocopy the effects of Ca^2+^ elevation induced by MPP^+^ and A23187.

Taken together, our results demonstrate that during oxidative stress, rise in cytosolic Ca^2+^ alone is not enough to augment cytotoxic mtROS production, but deployment of Zn^2+^ as a downstream signal is essential.

### Zn^2+^ inhibition of complexes I and III accounts for MPP^+^ (Ca^2+^)- induced mitochondrial ROS upregulation

We next asked how the MPP^+^-generated ionic signals exacerbate mtROS production. Although mitochondria have more than ten ROS generating sites, complexes I and III produce the majority (∼90%) of mtROS^1, 2, 3^. We therefore examined the roles of these two complexes using compounds that selectively suppress electron leak from complex I (S1QEL) ^31^ and complex III (S3QEL) ^32^. Both compounds attenuated MPP^+^-induced mtROS production (Figure 7A-B) as well as cell death (Figure 7C-D). However, S3QEL was more effective than S1QEL, even though its reported EC_50_ value in the ROS assay (∼1.75 µM) is more than one order of magnitude greater than that of S1QEL (∼ 0.06 µM)^31, 32^.

**Figure 7:**
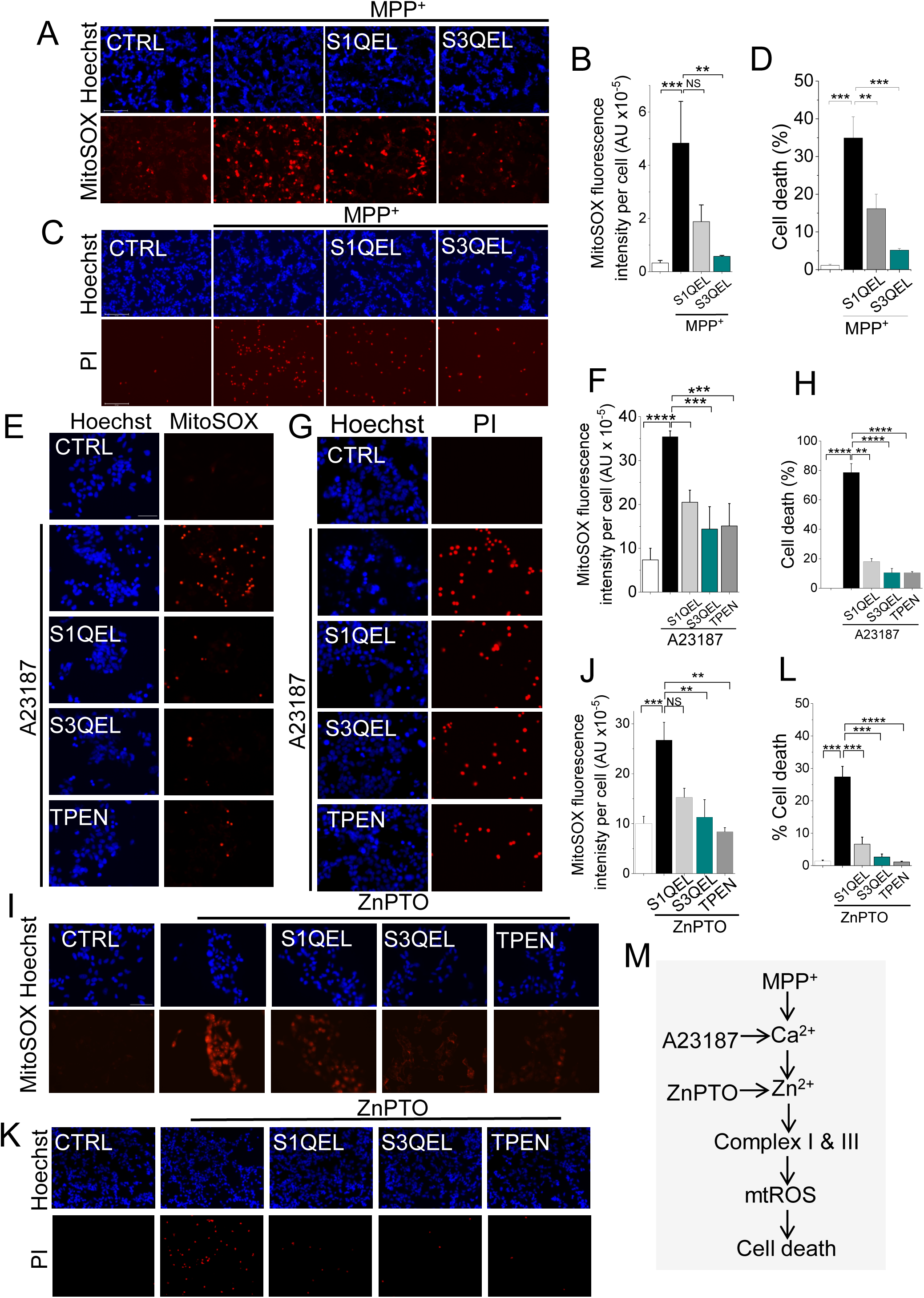
Zn^2+^ inhibits complexes I and III to generate cytotoxic levels of mitochondrial ROS. (**A-D**) MPP^+^-induced mtROS production and SH-SY5Y cell death are rescued by chemical suppressors of electron escape mainly from complex III. SH-SY5Y cells were pretreated with either S1QEL (10 µM) or S3QEL (5 µM) for 1 hour, and then treated MPP ^+^ (1 mM, 24 hours, 37°C). (**A-B**) S1QEL and S3QEL rescue MPP^+^-induced mtROS production. (**A**) Fluorescent images of cells stained for mtROS (Mito-SOX) and nuclei (Hoechst); scale bar = 125 µm. (**B**) Mean ± SEM of fluorescence intensity. (**C-D**) S1QEL and S3QEL rescue MPP^+^-induced cell death. (**C**) Fluorescent images of cells stained for cell death (Hoechst and PI); scale bar =125 µm. (**D**) Mean ± SEM of % PI positive dead cells. (**E-H**) A23187-induced ROS production and cell death are rescued by S1QEL and S3QEL as well as TPEN. S H-SY5Y cells were pretreated with S1QEL (10 µM) or S3QEL (5 µM) or TPEN (2 µM) for 1 hour, and then treated with 2 µM A23187 for 4 hours at 37°C. (**E**) Fluorescent images of cells stained for mtROS (Mito-SOX) and nuclei (Hoechst); scale bar = 75 µm. (**F**) Mean ± SEM of fluorescence intensity. (**G-H**) S1QEL and S3QEL rescue A23187-induced cell death. (**G**) Fluorescent images of cells stained for cell death (Hoechst and PI); scale bar =125 µm. (**H**) Mean ± SEM of % PI positive dead cells. (**I-L**) ZnPTO-induced ROS production and cell death are rescued by S1QEL and S3QEL as well as TPEN. Cells were pretreated as outlined in (E-H) and then exposed to ZnPTO (1 µM) for 2 hrs. (**I**) Fluorescent images of cells stained for mtROS (Mito-SOX) and nuclei (Hoechst); scale bar = 75 µm. (**J**) Mean ± SEM of fluorescence intensity. (**K-L**) S1QEL and S3QEL rescue ZnPTO-induced cell death. (**K**) Fluorescent images of cells stained for cell death (Hoechst and PI); scale bar =125 µm. (**L**) Mean ± SEM of % PI positive dead cells. Mean data in B, D, F, H, J and L are from three independent experiments (n =3); * *p* ˂ 0.05**; *p* ˂ 0.01; *** *p* ˂ 0.001; **** *p* ˂ 0.0001 from One-way Anova, post-hoc Tukey test. (**M**) Schematic summary of the findings.

We next substituted ionophores for MPP^+^ to determine the effects of Ca^2+^ and Zn^2+^ on the complexes. As with MPP^+^, both the QEL compounds suppressed the effects of A23187 on mtROS production (Figure 7E-F) and cell death (Figure 7G-H); these effects of QEL compounds are very much like TPEN (Figure 7E-H). Likewise, the QEL compounds were able to mitigate ZnPTO-induced mtROS production (Figure 7I-J) and cell death (Figure 7K-L). However, while S3QEL appeared to be slightly more effective than S1QEL, this difference did not reach statistical significance, presumably because ionophores raise the metal ion concentrations to levels well beyond what the pathological insults can achieve.

Collectively, these data suggest that MPP^+^ exacerbates mtROS production by generating Ca^2+^-driven Zn^2+^ signals that act on complexes I and III, primarily complex III (Figure 7M). Importantly, our data provide a fundamental new insight into how Ca^2+^ affects ROS production: while Ca^2+^ can increase ROS production due to its ability to increase in electron transport through the complexes ^10, 11^, for it to escalate ROS production to cytotoxic levels Zn^2+^ participation is mandatory.

## Discussion

Here we report the discovery of a signalling circuit that deciphers the complex mechanism underlying the production of pathogenic levels of mtROS. Although several signalling molecules including Ca^2+10, 11, 33^, mitochondrial complexes^1, 6, 7, 34, 35, 36^ and NOX2^13, 25, 37^ have been implicated, the question of whether and how they interact to drive mtROS production remained unclear. To address this, we have used a cellular model of PD^18^ wherein we exposed the SH-SY5Y cells to the PD causing MPP^+^ toxin and monitored the signalling events that upregulate mtROS production. The findings revealed that Ca^2+^, NOX2 and respiratory complexes form a signalling circuit by engaging TRPM2 channels and Zn^2+^ ions. Our study reveals several fundamental insights into the mechanism. First, MPP^+^ co-activates TRPM2 channels and NOX2 to rise cytoplasmic Ca^2+^. Second, Ca^2+^ targets mitochondrial complexes I and III to produce almost all pathogenic ROS; however, contrary to the widely held view^10, 11, 33^, the Ca^2+^ effect is not direct, but is mediated by Zn^2+^. Third, the signalling circuit plays no role in basal ROS production but is activated during external stress to drive ROS production to cytotoxic levels (depicted schematically in Figure 8).

**Figure 8.**
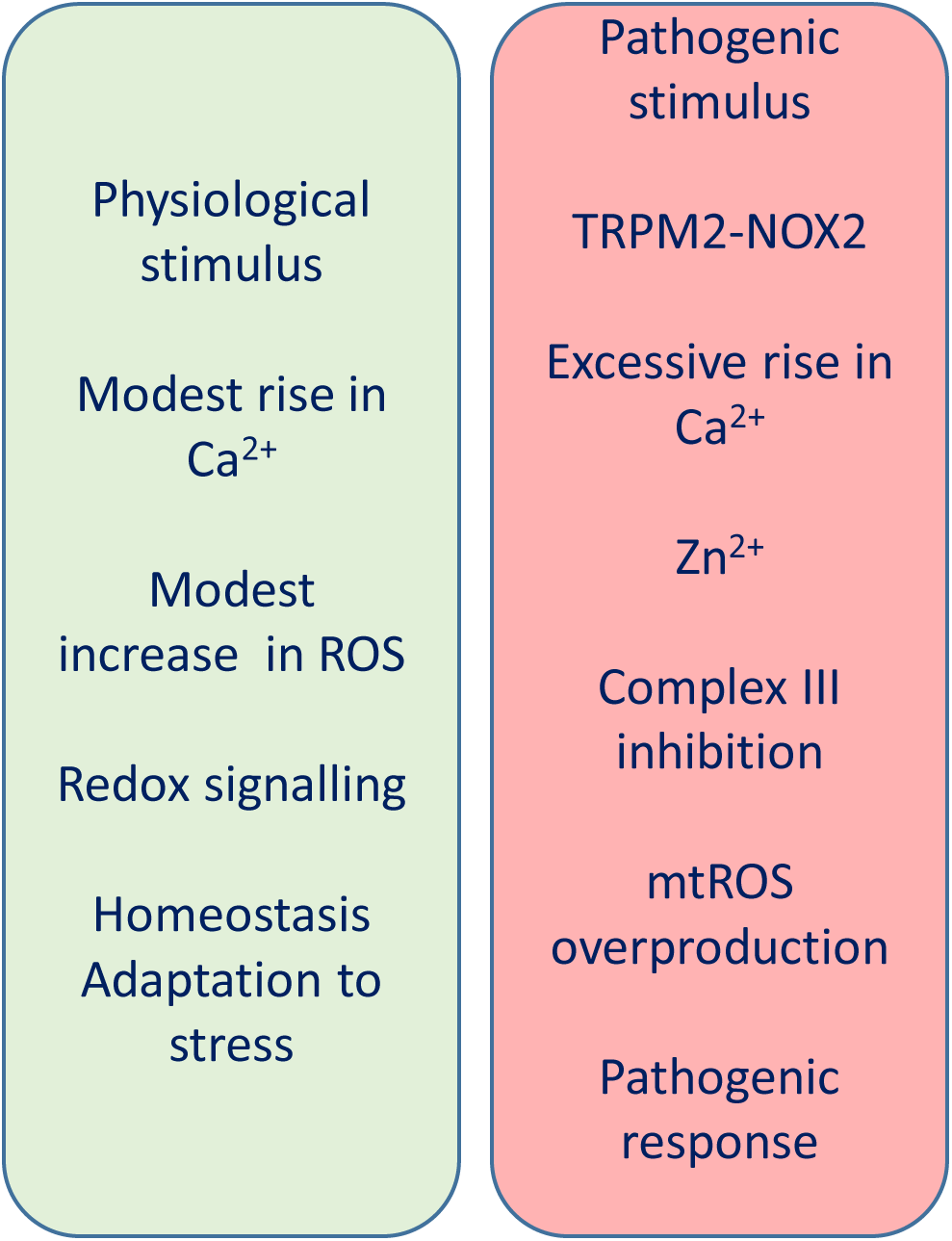
A district ROS generating signalling circuit (right panel) determines the pathological outcome. Unlike the mechanism responsible for the generation of ROS required for cell homeostasis and adaptation to stress (left panel), this signalling circuit involves activation of TRPM2-NOX2 duo to rise cytosolic Ca^2+^ to supra-physiological concentration to generate Zn^2+^ signals that, by inhibiting complex III, triggers excessive mitochondrial ROS production, leading to pathological outcome (right panel).

### MPP^+^ upregulates mitochondrial ROS production *indirectly* by generating Ca^2+^ and Zn^2+^ signals

Although MPP^+^ can increase cytoplasmic Ca^2+^ in SH-SY5Y cells^19^ (Figure 2A), Ca^2+^ has not been implicated in MPP^+^ induced ROS production. Instead, the current notion is that toxin inhibits complex I directly to stimulate ROS production ^38^. Although this view is based on studies with isolated mitochondria^24, 38^ and has been challenged by *in vivo* studies where knock-out of complex I failed to prevent dopaminergic neuronal loss^39^, the historic view continues to prevail. Given that Ca^2+^ is known to signal mtROS production^10, 11, 33^, we asked whether the ROS-inducing effect of MPP^+^ is, at least in part, owing to its ability to rise cytoplasmic Ca^2+^. Indeed, chelation of Ca^2+^ with BAPTA abolished the ability of MPP^+^ to augment cytotoxic mtROS production (Figure 2), suggesting that the MPP^+^ effect is almost entirely mediated by Ca^2+^.

However, BAPTA is not specific for Ca^2+^, but binds other metal ions, especially Zn^2+^ which has been shown to upregulate mtROS production in neuronal cells ^22^. Indeed, chelation of Zn^2+^ with TPEN, using a concentration (1 µM) too low to chelate Ca ^2+^, was as effective as BAPTA in mitigating MPP^+^ induced ROS production (Figure 2). Thus, we conclude that ability of MPP^+^ to trigger cytotoxic mtROS is not due to its direct effect on complex I, rather it is mediated by Ca^2+^ and Zn^2+^ ions.

### MPP^+^ induced Ca^2+^ influx requires simultaneous activation of TRPM2 channels and NADPH oxidase-2

We next asked how MPP^+^ increases cytoplasmic Ca^2+^. Although neuronal cells have multiple mechanisms to regulate Ca^2+^ ^11^, inhibition of Ca^2+^ influx through TRPM2 channels alone was sufficient to abolish MPP^+^-induced mtROS overproduction (Figure 3), indicating the unique role TRPM2 channels play in mtROS production. However, TRPM2 channels, paradoxically, require ROS for activation ^27, 40^. We demonstrate that ROS required for TRPM2 activation comes from NOX2^30^ activation (Figure 5). The finding that NOX2 inhibitors (Figure 5) are as effective as TRPM2 inhibitors (Figure 3-4) in mitigating MPP^+^-induced Ca^2+^ influx, and the subsequent mtROS production, raise the possibility that synchronised activation of NOX2 and TRPM2 likely occurs during MPP^+^ intoxication. Such mutual cooperation would allow NOX2 to provide ROS necessary for TRPM2 activation, and TRPM2 activation to supply Ca^2+^ for NOX2 activation^37^. Such functional interplay between TRPM2 and NOX2 potentially establishes a positive feedback loop near the plasma membrane to amplify the Ca^2+^ signals required for mtROS upregulation.

Although molecular basis for the interplay remains to be determined, it is interesting to note that upregulation of both NOX2 ^13, 37^ and TRPM2 channels^16, 17^ has been reported in numerous ROS-associated linked diseases. Furthermore, pharmacological, or genetic intervention of NOX2 ^13, 37^ and TRPM2 channels ^16, 17^ produce similar positive outcomes in experimental models of numerous ROS-linked diseases. Notably, activity of PARP1-an enzyme that produces ADPR -an endogenous activator of TRPM2 channels^15, 27, 40^- is markedly upregulated in several neurodegenerative diseases, including PD and Alzheimer’s disease ^41,42^. It therefore seems likely that co-activation of NOX2 and TRPM2 channels is a common mechanism shared by most ROS-linked diseases.

### For Ca^2+^ to induce mtROS overproduction requires Zn^2+^ mediation

We next asked how Zn^2+^ plays as an important role as Ca^2+^ in upregulating mtROS. To address this, we hypothesised a hierarchical relationship between the two signalling ions, with Zn^2+^ acting downstream of Ca^2+^. Consistent with this hypothesis, calcium ionophore-induced cytosolic Ca^2+^ rise induced mtROS production, yet this effect was entirely abrogated by TPEN (Figure 6). Moreover, elevation of intracellular Zn^2+^ with the ionophore, ZnPTO, led to mtROS production independently of Ca^2+^ (Figure 6). Collectively, these data argue that when cytoplasmic Ca^2+^ levels reach supraphysiological concentrations, Zn^2+^ acts downstream of Ca^2+^ to drive mtROS production to pathogenic levels.

Numerous reports have linked Zn^2+^ to PD pathology. Frist, Zn^2+^ is recognised as an environmental risk factor for PD ^43^. Second, accumulation of free Zn^2+^ is a key feature of degenerating dopaminergic neurons^44^ with a recent machine-learning study placing Zn^2+^ accumulation as a top predictor of PD^45^. Third, post-mortem examination of brains of PD patients^46^, Lewy body-injected macaque monkeys^45^ and MPP^+^ intoxicated mice ^44^ display deposits of labile Zn^2+^. Fourth, zinc transporters are upregulated in the brains of PD patients ^45^. Fifth, Zn^2+^ chelators protect mice against neurodegeneration in substantia nigra in *in vivo* models^47^. Finally, mutations in the PD associated *PARK9* gene disrupt Zn^2+^ as well as ROS homeostasis^48^. Thus, our findings provide a mechanistic explanation for the well-documented, but poorly understood role of Zn^2+^ in PD pathogenesis.

It is likely that in other ROS-driven diseases, where Ca^2+^ is implicated, Zn^2+^ plays a downstream role in generating pathogenic ROS. The beneficial effects of Zn^2+^ chelators in some experimental disease models including neurodegenerative^47, 49, 50^ and metabolic diseases^51^ lend some support to this plausibility.

### Zn^2+^ inhibition of complexes I and III accounts for MPP^+^ (Ca^2+^)- induced mitochondrial ROS upregulation

Guided by the previous reports that Zn^2+^ can inhibit complexes I and III ^52, 53^, we asked whether MPP^+^, via Zn^2+^, targets these complexes. Using compounds that selectively quench electron escape from complex I (S1QEL) and complex III (S3QEL) ^7, 32^, we demonstrate that MPP^+^ targets both complex I and III to induce mtROS production (Figure 7). These compounds suppressed MPP^+^-induced mtROS production, as well as mtROS induced by the calcium and zinc ionophores. These findings support the idea that the ability of MPP^+^ to induce excess mtROS stems from its ability to generate the Zn^2+^ signals that act on mitochondrial complexes. Our data showed that the MPP^+^ effect is more pronounced on complex III (Figure 7). This is presumably due to the much greater affinity of Zn^2+^ for complex III (Ki = 0.1 µM) ^52, 53^ compared with complex I (IC_50_10-50 µM) ^53^. This is consistent with the presence of a validated Zn^2+^ inhibitory binding residue, E within the catalytic centre of complex III, containing the highly conserved P**E**WY motif^54^. Complex I appears to lack Zn^2+^ sensitive residues within the catalytic centre, but Zn^2+^ was found in the regulatory 13 KDa accessory subunit of the complex^55^, however with no demonstrated role.

We consider the above findings in the context of pathophysiology. Under physiological conditions, the concentration of mitochondrial Zn^2+^ is too low (∼60 pM)^56^ to stimulate ROS from complexes I and III. However, basal ROS production continues to occur due to the ability of Ca^2+^ to stimulate electron flow and thereby increase ROS production from the complexes as well as other sites in mitochondria. During chronic stress, mitochondrial Zn^2+^ uptake increases, leading to the inhibition mostly of complex III due to its high affinity for Zn^2+^. From a pathogenic perspective, targeting complex III would seems advantageous because complex III releases O_2_^.-^ into the intermembrane space, from where it can more readily escape into the cytoplasm compared to the ROS released into the mitochondrial matrix from complex I ^57^.

Consistent with the dominant role of complex III, genetic depletion of the *Ndufs4* gene, which encodes a functional subunit of complex I, failed to prevent MPP^+^-induced ROS generation and death of dopamine-secreting neurons ^39^. Furthermore, a recent study reported a link between autosomal dominant mutations in *UQCRC1*, encoding a subunit of complex III, and familial Parkinson’s disease ^58^. These findings collectively suggest that the role of complex I in pathogenic mtROS production in PD, relative to complex III, may have been exaggerated in the literature.

### Conclusions and implications of the study

In summary, our findings present novel mechanistic insights into the upregulation of mtROS in a well-established cellular model of PD. Our work unravelled a signalling circuit encompassing NOX2, TRPM2, Ca^2+^, Zn^2+^ and complexes I/III. The presence of excessive NOX2 ^13^, Zn^2+^ ^59^, and oxidatively damaged biomolecules ^60^ in the post-mortem brains of PD patients, coupled with the genetic association between complex III and familial PD^58^, underscore the pathological relevance of this signalling circuit. Interestingly, toxic aggregates of α-synuclein have been shown to induce Ca^2+^ influx^61^, raising the possibility that the pathological effects of α-synuclein could be due to activation of the same signalling circuit. Although our findings were made using the cellular models of PD and diabetes, it is likely that this signalling circuit operates in a multitude of other conditions where mitochondrial dysfunction is a common denominator; these include other neurodegenerative diseases, metabolic diseases including diabetes^5, 6^, and perhaps ageing^62^. The fact that the signalling circuit is selectively activated by pathological insults suggests that it may represent a safe and effective therapeutic target for PD and a host of other diseases sharing common mechanisms.

## Methods

### Cell culture

SH-SY5Y cells (CRL-2266, Manassas, VA, USA) were cultured in DMEM-GlutaMAX^TM^-1 (31966-021, Thermo Fisher Scientific) supplemented with 10% foetal bovine serum (FBS), and penicillin (100 U/ml) and streptomycin (100 µg/ml) (P0781, Sigma-Aldrich) at 37°C in a humidified 5% CO_2_ incubator. Cells were grown to 70-80% confluency before passaging or plating out for experiments. HEK293-TRPM2^tet^ cells (kind gift from Dr A.M Scharenberg, University of Washington, Seattle, WA, USA) were cultured in the same medium, but contained additional selection antibiotics, Zeocin (400 µg/ml; P/N 46-0509, Invitrogen) and Blasticidin (5 µg/ml; ant-bl, InvivoGen). Antibiotics were absent in the media used for experiments. To induce TRPM2 expression, HEK293-TRPM2^tet^ cells were treated with tetracycline hydrochloride (1 µg/ml; T7660, Sigma-Aldrich) for 48 hours. INS-832/13 cell line (SCC207, Merck Millipore) was cultured in RPMI 1640 medium supplemented with 10% FBS, 100 U/ml penicillin, 100 µg/ml streptomycin, 10 mM HEPES, 2 mM L-glutamine, 1 mM sodium pyruvate, and 50 µM β-mercaptoethanol.

### Transfections

SH-SY5Y cells were transfected using Lipofectamine^®^ RNAiMAX according to the instructions of the manufacturer (56530, Invitrogen). Cells were grown in 24 -well plates to ∼60% confluency and transfected with small interfering RNA specific to TRPM2 (TRPM2-siRNA, 5′-GAAAGAAUGCGUGUAUUUUGUAA-3′, custom-made by Dharmacon) or scrambled control siRNA (Scr-siRNA: 4390846, Ambion) in Opti-MEM^TM^ using 25 nM siRNA and 1 ul of Lipofectamine^®^ RNAiMAX^28^. Experiments were performed 48 hours later.

### Treatments

Cells were plated (at 50 µg/ml) in poly-L-lysine (A003, Millipore) coated 96-well plates and grown in complete medium for 24 hours to ∼50% confluency. Cells were treated with the desired reagent prepared in the complete medium from stock solutions for the desired length of time at 37° C (see figure legends for specific details). Stock solutions were prepared as follows: 1-Methyl-4-phenylpyridinium (MPP^+^) iodide (D048, Sigma), (2-(2,2,6,6-Tetramethylpiperidin-1-oxyl-4-ylamino)-2-oxoethyl)triphenylphosphonium chloride, 2,2,6,6-Tetramethylpiperidin-1-yl)oxyl or (2,2,6,6-tetramethylpiperidin-1-yl)oxidanyl (TEMPO, A12497, AlfaAesar), Mito-TEMPO (SML0737, Sigma-Aldrich), N,N,N’,N’-tetrakis(2-pyridinylmethyl)-1,2-ethanediamine (TPEN, P4413, Sigma-Aldrich), 1,2-bis-(aminophenoxy)- ethane-N,N,N’,N’-tetra-acetic acid-acetoxymethyl ester (BAPTA-AM, P4758, ApexBio), (2- (Dimethylamino)-N-(6-oxo-5,6-dihydrophenanthridin-2-yl)acetamide hydrochloride (PJ34; A41159, ApexBio) N-(p-amylcinnamoyl) anthranilic acid (ACA, A8486, Sigma-Aldrich) A23187 (7522, Sigma-Aldrich), 1-Hydroxy-2-pyridinethione sodium salt (sodium pyrithione; H3216, Sigma-Aldrich), 4-hydroxy-3-methoxy-acetophenone (Apocynin, 73536, Merck), gp-91-ds-tat (AS-63818, AnaSpec; Fremont, CA, USA), 2’,7’-dichlorodihydrofluorescein diacetate (H_2_DCF-DA, D399, Invitrogen), dihydroethidine (DHE, 891IME, Bioserv), S1QEL 1.1 (SML1948, Sigma-Aldrich) and S3QEL 2 (SML1554, Sigma) were prepared in DMSO (BS-2245K, Bioserv). ZnPTO was prepared by mixing, in 1:3 ratio, aqueous solution of 1 mM zinc chloride (12973634, Fluka Chemical) with alcoholic solution of 3 mM sodium pyrithione. The medium in the wells was replaced with the treatment media and incubated at 37 °C for the desired length of time, as specified in figure legends.

### Cell viability assays

Following the treatment with the desired reagents, cells were washed with Hanks Balanced Salt solution (HBSS, 17420014, Corning) and stained with HBSS containing propidium iodide (5 µg/ml; P4170, Sigma-Aldrich) and Hoechst 33342 (4 µM, 5117, TORCIS Bioscience) for 30 minutes. Stained cells were imaged using an epifluorescent microscope (EVOS® FL Auto Imaging System; Thermo Scientific™ Invitrogen™). Percent cell death was calculated from the ratio of propidium iodide (PI) to Hoechst-stained cells.

### ROS detection

Total ROS was detected using H_2_DCF-DA or DHE, and mitochondrial ROS was detected using MitoSOX™ Red (M36008, Invitrogen). Following the desired treatments, media were replaced with HBSS containing 10 µM H_2_DCF-DA or 5 µM DHE or 5 µM MitoSOX™ Red and incubated for 30 min at 37°C. Cells were counter-stained with Hoechst 33342 and washed three times with HBSS, before imaging using the EVOS FL Auto 2 Imaging System fitted with a 20x or 40x objective and DAPI (excitation, 357 nm; emission, 470 nm for Hoechst), GFP (excitation, 470 nm; emission 525 nm for DCF) and RFP filter (525 nm excitation, 593 nm emission for DHE and MitoSOX™ Red) filters. In some experiments, LSM700 inverted confocal microscope fitted with 63x/1.4 NA oil objective (excitation, 500 nm; emission, 582 nm) was used.

### Ca^2+^ detection

Changes in intracellular Ca^2+^ were recorded using the ratiometric dye Fura-2-AM (F1201, Life Technologies). HEK293-TRPM2^tet^ cells grown in 96-well plates (Sarstedt) were preloaded with 1 µM Fura-2-AM/ 0.02% Pluronic® F127 (P-3000MP, ThermoFisher) in 100 ul HBSS. Changes in fluorescence due to rise in intracellular Ca ^2+^ were monitored using FlexStation^®^III (Molecular Devices). Cells were simultaneously excited at 340 nm and 380 nm and emission was recorded at 5s intervals at 510 nm to determine Ca ^2+^-bound and Ca^2+^-free Fura-2 respectively. The fluorescence ratio of 510 nm/340 nm to 510 nm/380 nm corresponds to the intracellular Ca^2+^ concentration. After taking basal readings, 20 µl of 15 mM H_2_O_2_ (22460250, Acros Organics) was injected into each well and the recordings continued.

To detect Ca^2+^ changes in SH-SY5Y cells, cells were grown in µ-Slide 8 well ^high^ ibiTreat (80806 Thistle Scientific) or 96-well plates to approximately 60% confluency and preloaded with 1 µM Fluo4-AM (F14201, Invitrogen) /0.02% Pluronic® F127 in HBSS. Following treatment with vehicle or test reagents at 37°C for the desired length of time, fluorescence was captured using the LSM700 inverted confocal microscope (excitation 494 nm; emission, 519 nm) or EVOS® FL Auto 2 Imaging System (using a GFP filter).

### Zn^2+^ detection

To detect intracellular free Zn^2+^, cells were grown in µ-Slide 8 well ^high^ ibiTreat slides to about 50% confluency. After the desired treatments, cells were loaded with 2 µM FluoZin3-AM (F24195, Invitrogen) in the presence of 0.02% Pluronic® F127 in HBSS to stain Zn^2+^. After washing (2x, 5 min each) with HBSS, images were captured using the LSM700 inverted confocal microscope (excitation: 494 nm; emission: 519 nm).

### Western blotting

HEK293-TRPM2^tet^ cell pellets solubilised in SDS sample buffer were run on 4-15% gradient SDS-PAGE gel. Separated protein bands were transferred from the gel onto a nitrocellulose membrane. Nonspecific sites on the membrane were blocked in a blocking buffer (5% skimmed milk/0.1% TWEEN20/Tris-buffered saline). TRPM2-FLAG protein was probed with mouse anti-FLAG®M2 antibodies (F1804, Sigma; primary, 1:5000 in blocking buffer) in conjunction with HRP-conjugated goat anti-mouse IgG (A16078, novusbio; secondary 1:10000 dilution in blocking buffer). Antibody bound bands were detected with Lumigen PS-Atto chemiluminescence reagent (PSA-100, Lumigen, Inc., Southfield, MI48033) and imaged using Syngene G:Box XX6 (Syngene).

### Data analysis and presentation

Fluorescence intensity (arbitrary units, A.U) of cells stained with various fluophores (DCF, DHE, MitoSOX, Fluo −4 and FluoZin-3) was estimated using Image J as described before^28^. All experiments were performed at least three times (n), each in duplicate wells. Images were acquired from three random fields of each well. Fluorescence intensity values were normalised to the number of cells determined by counting the Hoechst-counter stained nuclei. Data are presented in bar charts as mean ± S.E.M. Statistical significance was determined using the One-way ANOVA (Origin), followed by Tukey’s posthoc test; probability (*P*) values are indicated with *, **, *** and **** which correspond to values of 0.05, 0.01, 0.001, and 0.0001 respectively.

## Acknowledgements

We thank various institutions for PhD scholarships: Kuwait University, Kuwait (MA), The Ministry of Higher Education and Scientific Research, The State of Libya (HI and ES), University of Leeds, Leeds, UK (MA and FL), the Chinese Scholarship Council, PRC (FL).

