## supplemental file for "A distinct signalling circuit upregulates cytotoxic ROS production in a cellular model of Parkinson’s disease: key roles for TRPM2, Zn^2+^ and complex III"

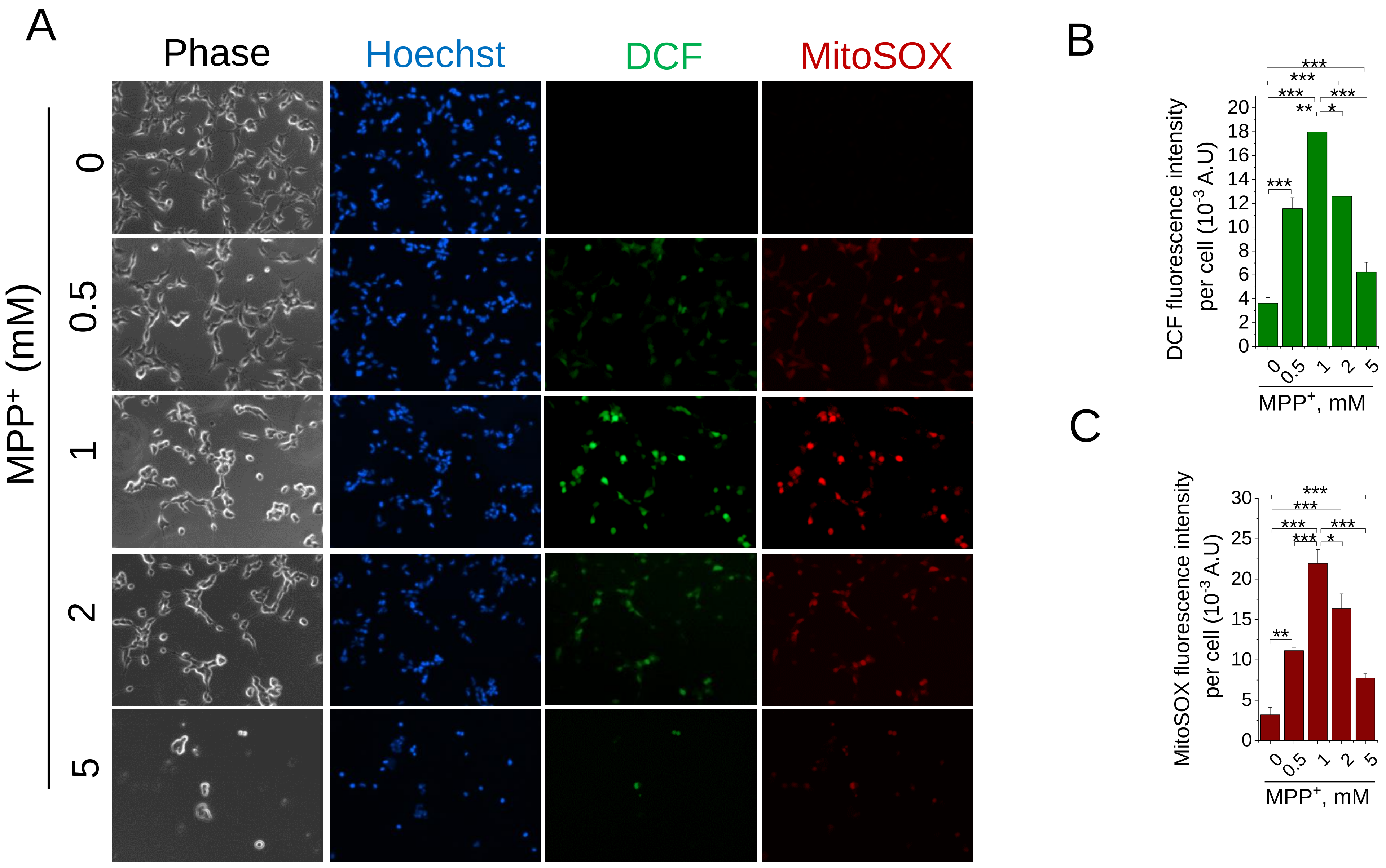


**Supplementary Figure 1**: **Dose dependent effect of MPP⁺ on total intracellular and mitochondrial ROS generation in SH-SY5Y cells.**

SH-SY5Y cells treated with the indicated concentrations of MPP⁺ for 24 h were stained for nuclei (Hoechst 33342), and for total (DCF) and mitochondrial (MitoSOX) ROS. **(A)** Representative phase and fluorescent images of stained cells. **(B-C)** the corresponding mean ± SEM of fluorescence intensity of DCF (B) and MitoSOX (C) from three independent experiments performed as in (A). ** $p < 0.01$; *** $p < 0.001$ from One-way Anova with post-hoc Tukey Test.
